# Structure of *C. elegans* TMC-2 complex suggests roles of lipid-mediated subunit contacts in mechanosensory transduction

**DOI:** 10.1101/2023.08.16.553618

**Authors:** Sarah Clark, Hanbin Jeong, Rich Posert, April Goehring, Eric Gouaux

## Abstract

Mechanotransduction is the process by which a mechanical force, such as touch, is converted into an electrical signal. Transmembrane channel-like (TMC) proteins are an evolutionarily-conserved family of ion channels whose function has been linked to a variety of mechanosensory processes, including hearing and balance sensation in vertebrates and locomotion in *Drosophila*. The molecular features that tune homologous TMC ion channel complexes to diverse mechanical stimuli are unknown. *Caenorhabditis elegans* express two TMC homologs, TMC-1 and TMC-2, both of which are the likely pore-forming subunits of mechanosensitive ion channels but differ in their expression pattern and functional role in the worm. Here we present the single particle cryo-electron microscopy structure of the native TMC-2 complex isolated from *C. elegans*. The complex is composed of two copies each of the pore-forming TMC-2 subunit, the calcium and integrin binding protein CALM-1 and the transmembrane inner ear protein TMIE. Comparison of the TMC-2 complex to the recently published cryo-EM structure of the *C. elegans* TMC-1 complex reveals differences in subunit composition and highlights conserved protein-lipid interactions, as well as other structural features, that together suggest a mechanism for TMC-mediated mechanosensory transduction.

**Significance Statement:** One mechanism by which organisms sense their environment is through the perception of mechanical stimuli such as sound, touch, and vibration. Transmembrane channel-like (TMC) proteins are ion channels whose function has been linked to a variety of mechanosensitive processes, including hearing and balance in vertebrates and touch sensation in worms. The molecular mechanisms by which TMCs respond to mechanical stimuli are unknown. Here we present the structure of the TMC-2 complex isolated from worms. Comparison of the TMC-2 complex to the recently solved structure of the worm TMC-1 complex highlights common structural features that are likely important for sensing mechanical stimuli yet also illuminates key differences that may explain the distinct functional roles of TMC-1 and TMC-2 in the worm.

## Introduction

Living organisms hear, sense touch, and perceive balance through mechanotransduction, the conversion of a mechanical force into an electrical signal. The primary sensors that mediate rapid responses to mechanical stimuli are mechanosensitive ion channels (1). Transmembrane channel-like (TMC) proteins are a family of ion channels whose function has been linked to a wide variety of mechanosensitive processes, including hearing in vertebrates (2, 3), water flow detection in zebrafish (4-6), and food texture sensation in *Drosophila* (7). Even within the same organism, TMC paralogs respond to different mechanical stimuli (8) and exhibit alternate responses to the same stimuli (2, 9). The features which tune homologous TMC proteins to diverse mechanical stimuli are unknown.

TMC proteins are conserved from *C. elegans* to humans *(10, 11)*. Vertebrates express eight TMC paralogs, of which TMC-1 and TMC-2 are important for the sensations of hearing and balance (11). TMC-1 and TMC-2 are the pore-forming subunits of the mechanoelectrical transduction complexes that are located in auditory and vestibular hair cells of the inner ear (12, 13). Although mammalian auditory hair cells express both TMC-1 and TMC-2, the expression patterns differ during development and they exhibit differences in current amplitude (2) and small molecule permeability (9). The six additional vertebrate TMC proteins (TMC-3 to TMC-8) are localized to various tissues, including the brain and colon (10, 11). Dysregulation of TMCs 4-8 have been linked to diverse human cancers (14, 15) and show promise as biomarkers for cancer prognosis and potential therapeutic targets (15). However, the function of TMCs 3-8 remains unknown.

Invertebrates also express orthologs of TMC proteins, many of which have been implicated in mechanosensory processes. *Drosophila* express a single TMC ortholog that is important for larval locomotion and food texture sensation (7, 16). The *C. elegans* genome encodes two *tmc* genes, *tmc-1* and *tmc-2*, both of which encode proteins that are subunits of mechanosensitive ion channel complexes (8). The *C. elegans* TMC-1 protein has been implicated in a plethora of cellular processes, including sodium chemosensation (17), alkaline sensation (18), modulation of membrane excitability (19), touch sensation (8), metabolism (20), and regulation of GABA signaling (21). TMC-1 is localized to mechanosensory and chemosensory neurons, as well as body wall and vulval muscle (8). By contrast, *C. elegans* TMC-2 is localized exclusively to body wall and vulval muscles (8) and its role in worm physiology is less well explored. We recently solved the structure of the *C. elegans* TMC-1 complex, illuminating structural elements that are likely important for mechanosensation and providing insight into the structure of the vertebrate mechanosensory transduction (MT) complexes (22). Given the differences in function and expression pattern of *C. elegans* TMC-1 and TMC-2, we were curious how the architecture of the TMC-2 complex differs from that of TMC-1.

Here we use cryo-electron microscopy (cryo-EM) to solve the structure of the native TMC-2 complex isolated from transgenic worms. Furthermore, to complement the structures of the TMC-1 and TMC-2 complexes, we used mass spectrometry to investigate whether additional proteins, such as ankyrin or UNC-44 in worms, are part of the TMC-2 complex. Previous research suggested that ankyrin couples TMC proteins to actin, functioning as an intracellular tether to relay mechanical force to the ion channel (8). The structure of the TMC-2 complex is strikingly similar to the TMC-1 complex, highlighting conserved structural features and protein-lipid interactions that shed light on how TMC complexes sense force. Further, the TMC-1 and TMC-2 complexes exhibit several key differences in the subunit composition and pore-lining amino acids that may render the complexes functionally distinct.

## Results

### Subunit composition and overall architecture of the TMC-2 complex

To isolate the native TMC-2 complex from *C. elegans*, we employed the same strategy that was used for the TMC-1 complex (22, 23). In brief, we generated a transgenic worm line wherein the 3’ end of the TMC-2 coding sequence was tagged with a nucleic acid sequence encoding a mVenus fluorophore and a 3xFLAG tag (*tmc2::mVenus*). Using spectral confocal imaging, we characterized the *tmc2::mVenus* worm line and demonstrated that the TMC-2-mVenus protein is expressed in the body wall and vulval muscles (Figure S1a), in agreement with previous studies (8). The expression level of TMC-2 in *C. elegans* is roughly 3-fold higher than that of TMC-1, as assessed by fluorescent-detection size-exclusion chromatography (FSEC) experiments (24) using an equivalent number of *tmc1::mVenus* and *tmc2::mVenus* worms. To isolate the TMC-2 complex, we homogenized ∼ 4 × 10^6^ *tmc2::mVenus* worms and performed affinity chromatography followed by fluorescent-detection size-exclusion chromatography (FSEC) (Figure S1b). The estimated molecular weight of the TMC-2 complex by FSEC was ∼750 kDa, slightly smaller than that of the TMC-1 complex, but far larger than the predicted 340 kDa molecular weight of a TMC-2 dimer, suggesting that auxiliary proteins are bound to TMC-2.

Mass spectrometry analysis of the TMC-2 complex revealed a similar complex composition to TMC-1 (22), with one notable difference. The auxiliary proteins calcium and integrin binding CALM-1 and transmembrane inner ear protein TMIE were both identified as subunits of the TMC-2 complex by mass spectrometry (Figure S1c), in accordance with mass spectrometry results of the TMC-1 complex. Intriguingly, however, the TMC-1 auxiliary protein arrestin (ARRD-6), was not identified by mass spectrometry or by cryo-EM analysis of the TMC-2 complex. Because ARRD-6 interacts exclusively with CALM-1 and the intracellular leaflet of the membrane, and does not directly interact with TMC-1, it is unclear why ARRD-6 did not co-purify with the TMC-2 complex. Differences in tissue localization between TMC-1 and TMC-2, specifically the absence of TMC-2 in sensory neurons, may explain the lack of TMC-2 association with ARRD-6. These results therefore suggest ARRD-6 may play a role in TMC-1-mediated mechanosensation in worm sensory neurons. Further, we did not find evidence for TMC-2 association with UNC-44, the worm ortholog of mammalian ankyrin. UNC-44 has been reported to interact with CALM-1 and play a vital role in TMC-1- and TMC-2-mediated mechanosensation in worms (8), yet we did not detect UNC-44 via mass spectrometry or cryo-EM in either the TMC-1 (22) or TMC-2 complexes. Likewise, we found no evidence for the presence of PEZO-1, the worm homolog of PIEZO-1 and 2, thus demonstrating that, at least in worms, TMC-1 or 2 do not form a stable complex with PEZO-1 (25).

To elucidate the architecture of the native TMC-2 complex, we performed single particle cryo-electron microscopy (cryo-EM) (Figure S2, Table S1). Approximately 100 ng of protein was isolated from *tmc2::mVenus* worms and placed on 2 nm carbon grids that had been glow discharged in the presence of amyl amine. Rounds of 2D classification of the selected particles revealed a high degree of heterogeneity that was not observed in the TMC-1 dataset, despite nearly identical sample and grid preparation methods for TMC-2, including the use of the same detergent as employed in the isolation of TMC-1 (22). The majority of TMC-2 particles were monomeric and exhibited a preferential ‘top-down’ orientation. We also observed an apparent ‘trans-dimer’ conformation that we discarded in 2D classification (Figure S2b). Particles that corresponded to a *cis*-dimeric complex were selected for subsequent refinement and 3D reconstruction, yielding a density map at 3.2 Å resolution, sufficient for model building (Figure 1, Figure S3). Although we attempted to reconstruct the monomeric and ‘trans-dimer’ conformations of the TMC-2 complex, we were not successful. This may be due to a combination of small protein size, possible conformational heterogeneity, and low particle numbers.

**Figure 1:**
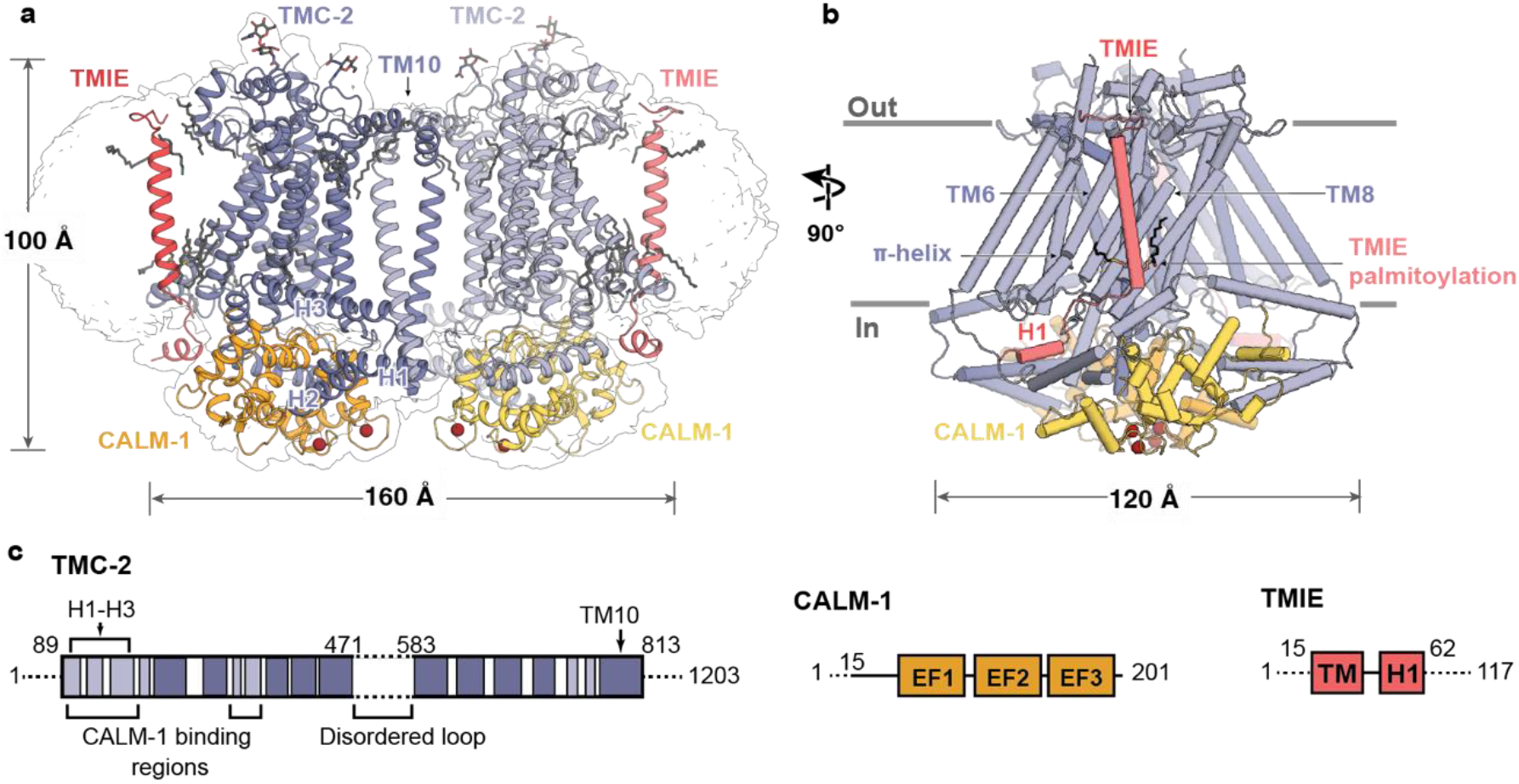
Architecture of the TMC-2 complex. **a**, Overall architecture of the TMC-2 complex, viewed parallel to the membrane. TMC-2 (dark and light purple), CALM-1 (orange and yellow), and TMIE (dark and light pink), are shown in a cartoon diagram. Lipid-like molecules and *N*-glycans are colored grey, while calcium ions red. The map refined without a mask is shown as a transparent envelope. **b**, A 90° rotated-view of the TMC-2 complex, viewed parallel to the membrane. Structural features of interest are indicated. α-Helices are shown as cylinders. **c**, Schematic representation of proteins that form the TMC-2 complex. For TMC-2, helices are shown in light purple and transmembrane helices are shown in dark purple. Dashed lines indicate regions that were not observed in the cryo-EM structure. H = helix, TM = transmembrane domain, EF1-EF3 = EF-hand domains.

The overall architecture of the TMC-2 complex is similar to TMC-1 (Figure 2). Although TMC-1 and TMC-2 share only 36% sequence identity, superposition of a single protomer from each structure yields a backbone root mean square deviation (RMSD) of 0.9 Å. As in TMC-1, the TMC-2 complex harbors two copies each of TMC-2, CALM-1, and TMIE (Figure 1a). Each TMC-2 protomer consists of ten transmembrane helices and the dimer interface is composed of domain-swapped transmembrane helix 10 (TM10). The cytosolic domain includes six helices oriented nearly parallel to the membrane plane, all of which form extensive interactions with CALM-1. The transmembrane domain exhibits the highest local resolution (Figure S3) and is decorated with an abundance of lipids or lipid-like molecules. Density features consistent with glycosylation are observed at N225, located in the loop between TM1 and TM2, as well as N748, located in the loop between TM9 and TM10. Peptides from both the 125-residue loop between TM5 and TM6 and the approximately 400-residue C-terminus were detected by mass spectrometry, but these regions were not visible in our map (Figure 1c). We suspect that this region is not visible due to heterogeneity, as we observed with TMC-1.

**Figure 2:**
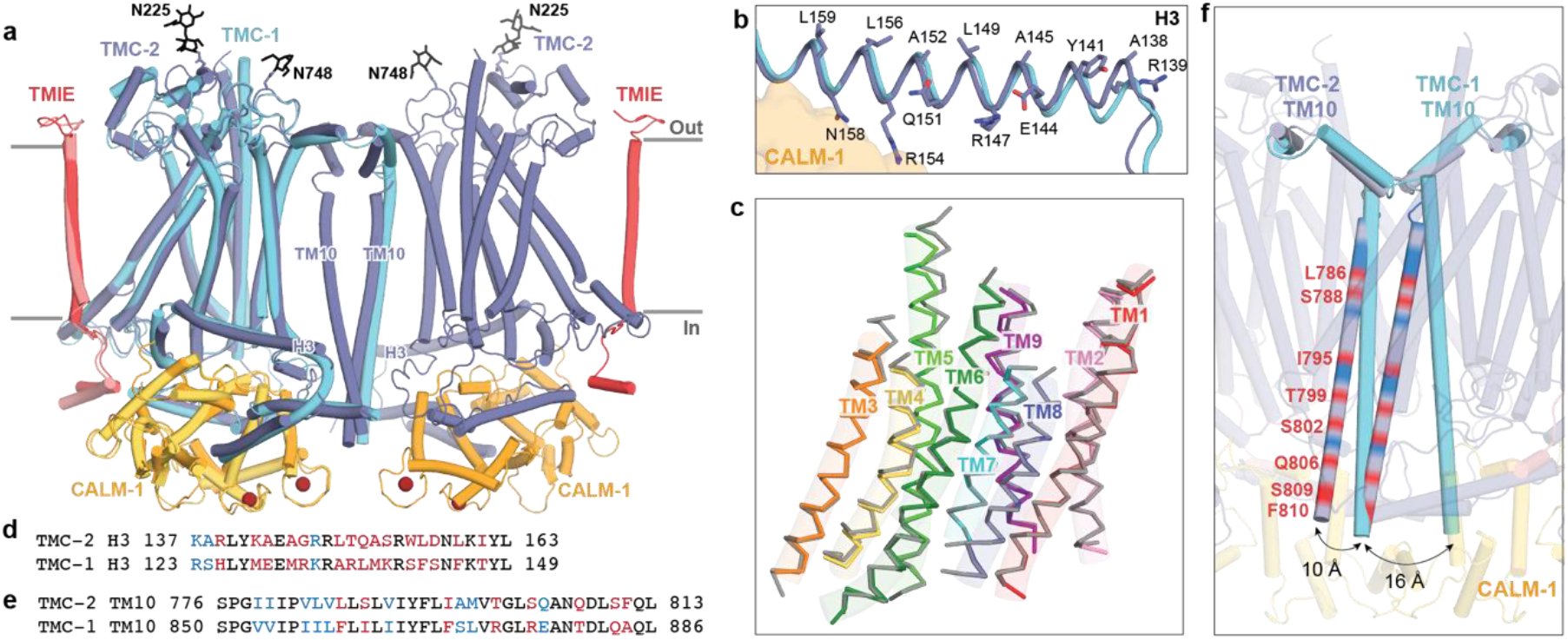
The TMC-1 and TMC-2 complexes have similar architectures, yet adopt distinct conformations at the dimer interface. **a**, Superposition of backbone atoms of the TMC-2 complex with backbone atoms of one protomer of the TMC-1 complex, shown on the left. TMC-1 is colored cyan and the CALM-1 and TMIE auxiliary subunits from the TMC-1 complex are colored yellow and pink, respectively. The TMC-2 complex is colored as in Figure 1. *N*-glycan molecules are colored black and calcium ions are colored dark red. **b**, TMC-2 helix H3, shown in purple, is superposed with TMC-1 helix H3, shown in cyan. Residues that confer an amphipathic nature upon the TMC-2 H3 helix are shown as sticks. **c**, The TMC-1 transmembrane helices (TM1-TM9), shown in grey, superposed on the TMC-2 transmembrane helices, shown in different colors, using backbone carbon atoms. **d**, Sequence alignment of the amino acid residues of the TMC-2 and TMC-1 H3 helices. Residues are colored black for identical, blue for conservatively substituted, or red for non-conservatively substituted. **e**, Sequence alignment of amino acid residues of the TMC-2 and TMC-1 TM10 regions. Residues are colored as in (d). **f**, Closeup view of the dimer interface of the superimposed complex from panel (a), rotated ∼45° about an axis perpendicular to the membrane plane, highlighting the different conformations of the TM10 regions in the two complexes. The positions of the non-conserved residues in TM10 of TMC-2, compared to TMC-1 and as defined in panel (e), are shown as red bands.

### TMC-1 and TMC-2 are structurally conserved aside from the dimer interface

Alignment of the TMC-1 and TMC-2 complex structures based on backbone atoms of a single TMC-1 or TMC-2 protomer reveals a high degree of structural conservation in all regions of the complex, including the transmembrane domain, cytoplasmic helices, and auxiliary protein positions (Figure 2a). The side chain conformations of conserved amino acids in the TMC-1 and TMC-2 transmembrane helices are remarkably similar, although several non-conservative substitutions are apparent in the cryo-EM density map that allow us to easily distinguish between the density maps and their corresponding structures (Figure S4).

Close comparison of the TMC-1 and TMC-2 complexes, however, reveals several notable differences. On the extracellular side of the complexes, all of the short loops and helices that link transmembrane domains are conserved at a sequence and structural level, aside from the long, unstructured loop that connects TM5 and TM6. This loop, which was not visible in either the TMC-1 or TMC-2 maps, is 75 residues shorter in TMC-2 and only exhibits 14% sequence identity to the TMC-1 loop. Further, while TMC-1 and TMC-2 are both glycosylated at N209 and N225, respectively, we observed a second density feature at a predicted glycosylation site in the TMC-2 map, located on N748 between TM8 and TM9 (Figure 2a). TMC-1 bears an arginine residue at this position (R821) and the surrounding four residues are also different in amino acid sequence.

The six cytoplasmic helices, all of which interact with the auxiliary protein CALM-1, are structurally conserved between TMC-1 and TMC-2. The helices that share the greatest degree of sequence identity are also those that form the majority of the interactions with CALM-1: H1, H2, H4, and H6. Helix H3, by contrast, only shares 33% sequence identity between TMC-1 and TMC-2 (Figure 2d). Despite the lack of sequence conservation, helix H3 retains the amphipathic nature that was observed in the TMC-1 structure (Figure 2b). Hydrophobic residues on the membrane-proximal side of H3 allow it to interact with the inner leaflet of the membrane bilayer, while polar and charged residues occupy the cytoplasmic side, several of which interact with CALM-1 (R154, N158). Amphipathic, cytosolic helices that are oriented parallel to the membrane bilayer have been observed in the structures of other mechanically-activated channels, such as OSCA2.1 (26) and PIEZO(27), and have been proposed to be a mechanistically important feature of mechanosensitive ion channels (1).

Superposition of the backbone atoms of TM1-TM9 from TMC-1 and TMC-2 reveals nearly identical positions for the transmembrane helices (Figure 2c) and sequence identity among the TMs ranges from 40-86%. However, TM10, which mediates the domain-swapped dimer interfaces, is structurally dissimilar in the TMC-1 and TMC-2 complexes (Figure 2e, f). Unlike TMC-1, TMC-2 is not C2 symmetric owing to the orientation of the TM10 helices, which are offset by 10-16 Å relative to the TMC-1 TM10 helices (Figure 2f). TM10 exhibits 52% sequence identity between TMC-1 and TMC-2 (Figure 2e), but the side chains of the non-conserved amino acids are directed away from the dimer interface. Dimerization of TMC-2 results in burial of 1,613 A^2^ of solvent accessible surface area, slightly less than the 1,781 A^2^ of surface area that is buried upon TMC-1 dimer formation, suggesting that fewer contacts exist to stabilize the TMC-2 dimer. Notably, the ∼400 C-terminal residues that follow TM10, which were not visible in the cryo-EM map, only share 22% sequence identity with TMC-1. It is possible that differences in the structure or flexibility of the TMC-1 and TMC-2 C-termini may also indirectly impact the conformation of TM10.

### Auxiliary protein binding interfaces and lipid-mediated interactions are conserved between TMC-1 and TMC-2

Mass spectrometry results indicate that *C. elegans* TMC-2 co-purifies with the auxiliary proteins CALM-1 and TMIE (Figure S1), both of which are predicted components of the vertebrate MT complex (5, 28, 29). In vertebrates, TMIE is known to play an important role in ion channel gating (30), while the specific role of CALM-1 in mechanosensory transduction is less well understood. Mechanotransduction currents are abolished in mice with null mutations in either TMIE and CALM-1, resulting in deafness (28, 29). The *C. elegans* TMC-1 and TMC-2 complexes share nearly identical binding interfaces for CALM-1 and TMIE, emphasizing the importance of these auxiliary protein interactions for MT complex function (Figure 3a-e). CALM-1 interacts with all six TMC-2 cytoplasmic helices through an extensive network of hydrogen bonds, electrostatic interactions, and hydrophobic contacts. The short loop that connects TMC-2 H1 and H2 harbors an abundance of basic residues that interact with an acidic patch on CALM-1 (Figure 3c) and TMC-2 H4-H6 are bound to CALM-1 through a mixture of polar and nonpolar interactions (Figure 3d). Nearly all of the TMC-1 and TMC-2 residues that mediate interaction with CALM-1 are conserved (Figure 3e), highlighting the high degree of conservation in the CALM-1 binding interface. Likewise, the regions of TMC-1 and TMC-2 that mediate interaction with TMIE are conserved, both in structure and in sequence (Figure 3b, f). Arginine residues in the cytoplasmic TMIE ‘elbow’ form hydrogen bonds with backbone atoms of TMC-2 TM6 and TM8, and TMIE R23 and W25 residues on the extracellular side interact with the loop between TMC-2 TM1 and TM2. TMIE also makes lipid-mediated hydrophobic contacts with the TMC-2 transmembrane domains through phospholipid molecules and palmitoylation of two TMIE cysteine residues. These lipid features and contact sites are conserved and are discussed in the following sections.

**Figure 3:**
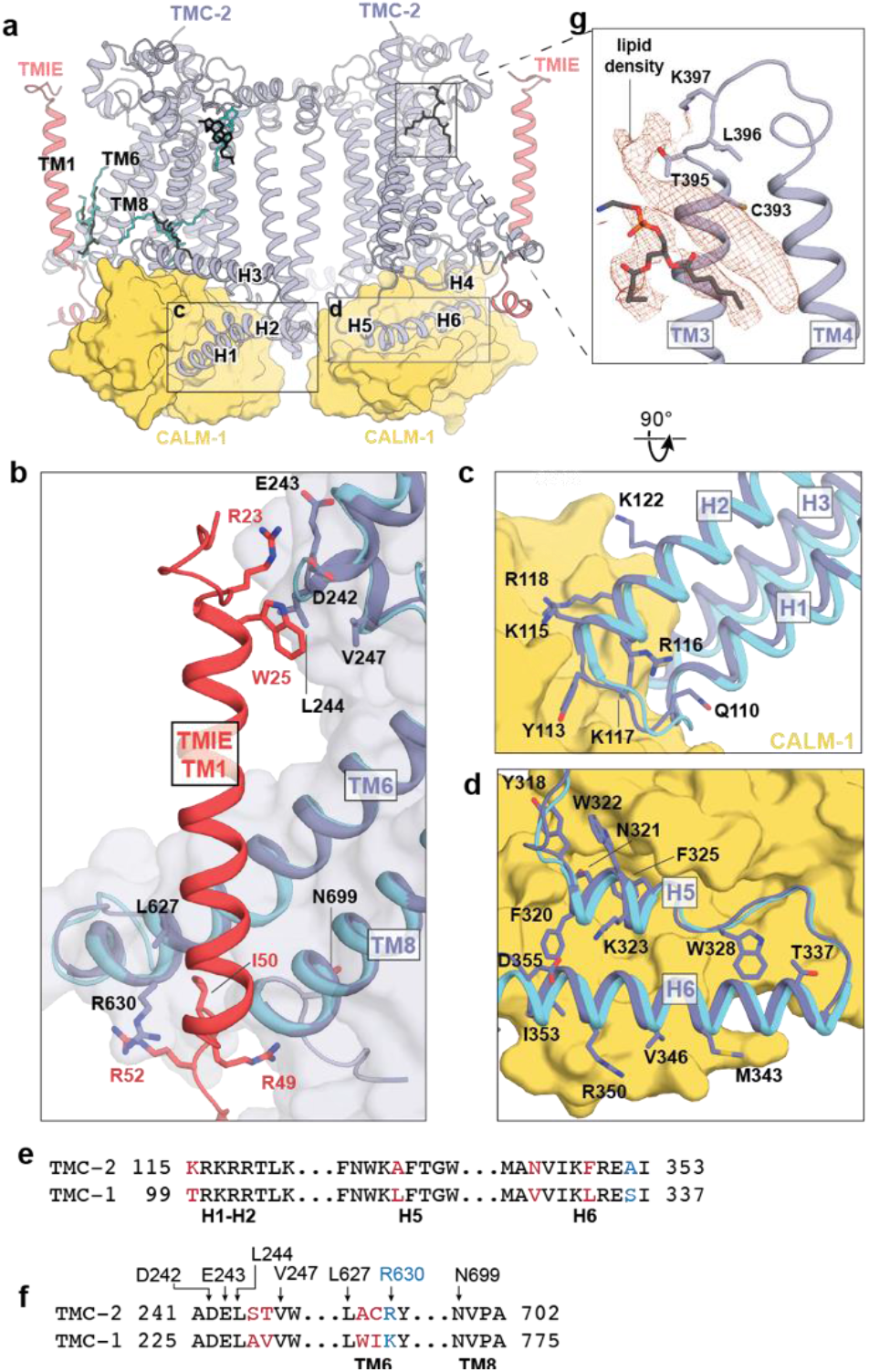
TMC-1 and TMC-2 share a conserved interaction interface with the auxiliary subunits TMIE and CALM-1. **a**, Lipids with strong density features and similar locations in the TMC-1 and TMC-2 density maps are shown in the context of the TMC-2 complex. TMC-1 lipids are cyan and TMC-2 lipids are black. **b**, The subunit interface of TMIE in the TMC-1 or TMC-2 complex. TMC-2 is purple, TMC-1 is cyan, and TMIE is red. Interacting residues are shown as sticks. **c**, The subunit interface of CALM-1 with TMC-1 or TMC-2 helices H1-H2. CALM-1 is colored yellow and interacting residues are shown as sticks. **d**, The subunit interface of CALM-1 with TMC-1 or TMC-2 helices H5-H6, colored as in (c). **e**, Sequence alignment of the residues from the cytoplasmic helices of TMC-1 and TMC-2 that form the interaction interface with CALM-1. Residues in black are conserved. Residues in blue are conservatively substituted and residues in red are not conserved. **f**, Sequence alignment of TMC-2 and TMC-1 residues that interact with TMIE. Residues are colored as in (e). **g**, Lipid-like density features observed in the TMC-2 map that were not observed in the TMC-1 map. Density map is shown in red and TMC-2 is shown in purple.

Multiple ordered lipids and lipid-like molecules decorate the TMC-1 and TMC-2 transmembrane domains (Figure 1a). Remarkably, many of the lipids with the strongest density features are found in the same locations in both structures (Figure 3a), suggesting that they play a role in complex stability or ion channel function, or both. A phospholipid occupies the cleft between TMC-2 H3 and TM1, and another resides in the cavity between the TMIE transmembrane helix and TMC-2 TM6 and TM8. The location of the ‘bridge’ lipid between the TMIE and TMC-1/2 transmembrane helices is of special interest because it makes hydrophobic contacts between structural elements that are predicted to be involved in ion channel gating. An additional density feature that is consistent with a cholesterol molecule resides proximal to the extracellular side of TM2 in both structures. Interestingly, we observed a prominent lipid-like density in the TMC-2 map that is not present in TMC-1 (Figure 3g). The lipid-like feature resides in the extracellular leaflet, occupying a crevice between TM3 and TM4. Further analysis, and possibly higher resolution reconstructions, will be required to determine the molecular composition of this lipid-like density feature.

### The putative pore of TMC-2 is distinct from TMC-1

The putative ion conduction pathway of TMC-2 follows the same trajectory as the conduction pathways in TMC-1 and the evolutionarily-related proteins TMEM16A (31, 32) and OSCA1.2 (26, 33) (Figure 4a). Like the structure of TMC-1, the ion conduction pore is lined by TM4-8. The pore appears closed and is blocked by three sites of constriction (Figure 4b, c). The extracellular pore entrance, which is lined by a mixture of polar and nonpolar residues, is followed by a narrow ‘neck’ region that is dominated by hydrophobic residues (Figure 4d). The remainder of the pore, towards the cytoplasm, is lined by polar and charged residues (Figure 4b, d).

**Figure 4:**
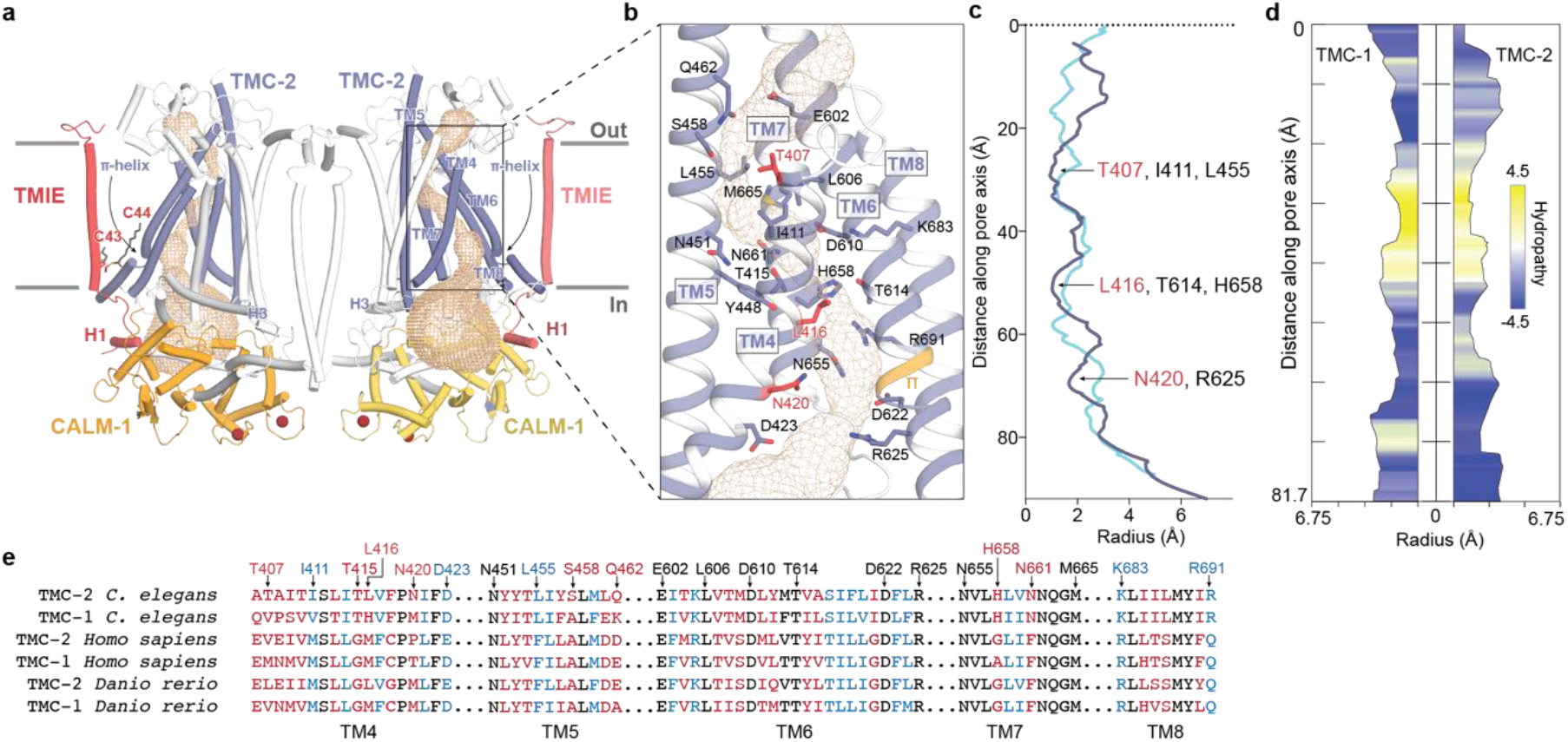
The putative ion conduction pore of TMC-2. **a**, The putative location of the pore is shown as gold mesh. The pore-lining helices, TM4-8, are shown in purple and the remaining TM helices are shown in white. **b**, An expanded view of the possible ion permeation pathway, showing pore-lining residues as sticks. Residues that are not conserved between TMC-1 and TMC-2 are shown in red. A π-helical turn within TM6 of TMC-2 is shown in orange. **c**, Radii of the TMC-1 and TMC-2 pores plotted along the pore axis, calculated by Mole 2.5. TMC-1 is shown in blue and TMC-2 is shown in purple. **d**, Hydropathy plot of the TMC-1 pore (left) and TMC-2 pore (right) calculated by Mole 2.5. Regions of the pore with high hydrophobicity values are colored yellow and regions with high hydrophilicity values are colored blue. The size of the pore is plotted along the pore axis. **e**, Multiple sequence alignment of selected residues from the putative TMC-2 and TMC-1 pore-lining helices of different species, with the residues colored as in panel (b).

The ion selectivity and permeation properties of the *C. elegans* TMC-1 and TMC-2 ion channels are not known, in large part due to an inability to express functional channels in a heterologous system. Studies of vertebrate MT channels in hair cells have demonstrated that they are cation-selective ion channels (34) with a high permeability for calcium (35). Analogous to *C. elegans* TMC-1, the composition of *C. elegans* TMC-2 pore-lining residues is not consistent with a cation-selective channel, which usually harbors a predominance of acidic residues. While four highly conserved acidic residues line the TMC-2 pore, there are also four basic residues scattered throughout the putative conduction pathway, two of which are conserved across multiple species (Figure 4b, e). It is unclear if these residues line the pore in its open conformation.

The majority of the pore-lining residues are conserved between TMC-1 and TMC-2, nevertheless. Intriguingly, TM4 contains three of the five residues that are not conserved. These amino acid differences all involve substitution of a polar or charged residue for a nonpolar residue. Mutagenesis studies of vertebrate TMC-1 in hair cells have demonstrated that TM4 and TM6 are important for mechanical gating of TMC-1 (36), suggesting that the aforementioned non-conservative substitutions in TM4 may alter the mechanical activation of *C. elegans* TMC-2 relative to TMC-1.

### Palmitoylation of TMIE suggests a mechanism for TMC-1 gating

Studies in vertebrate hair cells have demonstrated that TMIE alters the gating properties, unitary conductance, and ion selectivity of the vertebrate MT complex (30). The structure of *C. elegans* TMC-1 complex revealed that TMIE C44 is palmitoylated and the acyl chain extends towards TMC-1 pore-forming transmembrane helix TM8, potentially linking TMIE to the ion conduction pathway. We observed a second TMIE palmitoylation site on C43 in the TMC-2 complex structure that may have additional implications for gating of TMC ion channels (Figure 5a). Prompted by the palmitoylation of TMIE in the TMC-2 complex, we re-inspected the cryo-EM density maps of the TMC-1 complex and found evidence that TMIE C43 is also palmitoylated (Figure 5b). The palmitoyl groups of C43 and C44 are oriented almost identically in both the TMC-1 and TMC-2 complexes; the acyl chain of C43 extends in the direction of TM6 and the acyl chain of C44 makes hydrophobic contacts with nonpolar residues in TM8 (Figure 5). Interestingly, the palmitoyl groups of TMIE exhibit a conformation in which their acyl groups embrace the conserved ‘bridge’ lipid. While the density for a palmitoyl group on C43 in both the TMC-1 and TMC-2 reconstructions is weaker than the corresponding density associated with C44, we were nevertheless able to confidently fit a palmitoyl moiety to the density. Because the density for the palmitoyl group of C44 is stronger, we speculate that the two palmitoyl groups differ in their conformational mobility. How these differences are related to the function of the complex remains to be determined.

**Figure 5:**
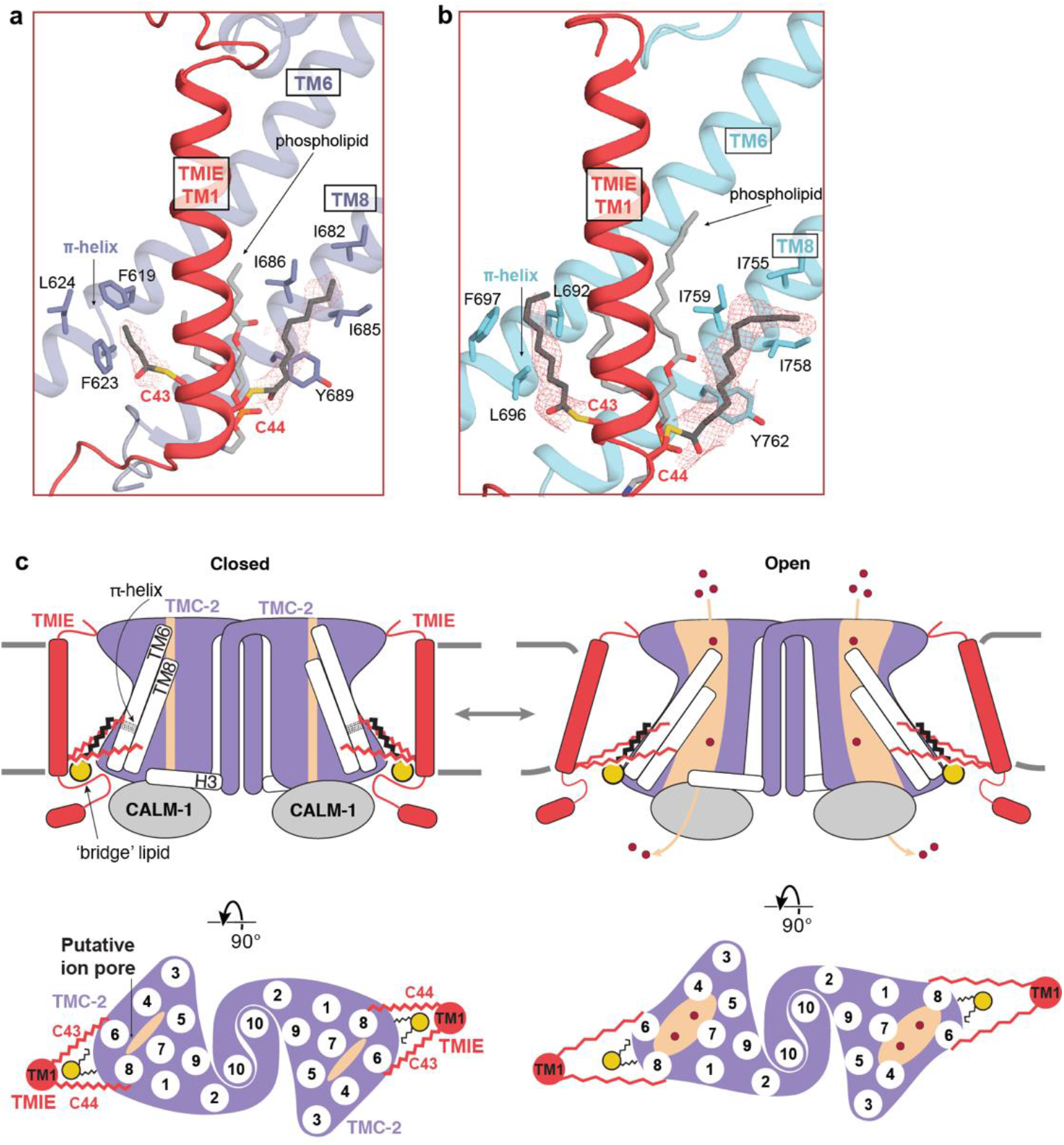
Lipid-mediated interactions between TMIE and TMC-1/2. **a**, Palmitoylation of TMIE C43 and C44 is observed in the TMC-2 complex. Interacting residues are shown as sticks and a phospholipid is shown in light grey. Cryo-EM density is shown as red mesh, at a contour level of 5.68 RMSD. **b**, Palmitoylation of TMIE C43 and C44 is also observed in the TMC-1 complex. TMC-1 is shown in cyan and other coloring is depicted as in (a). The density map contour level is 4.62 RMSD. **c**, Schematic illustrating how lipid-mediated interactions between TMIE and the TMC-1/2 transmembrane helices may translate forces to ion channel gating. Lipid interactions may stabilize the closed conformation of the channel (left). Disruption of the membrane due to mechanical forces could alter these lipid interactions, leading to ion channel opening (right). Phospholipid head groups are colored yellow.

Notably, the C43 palmitoyl group contacts two phenylalanine residues, F619 and F623, that are positioned on a π-helical turn of TMC-2 TM6. The π-helical turn is conserved in the TMC-1 structure, but the phenylalanine residues are replaced by leucine residues, L692 and L696 (Figure 5b). TMC-1 F697 further bolsters the hydrophobic interaction with the palmitoyl group. A leucine or phenylalanine residue is conserved in all three positions across multiple species (Figure 4e), indicating that these residues may be important for the indirect interaction between TMIE and TMC-1/2. As TMC-1/2 TM6 plays a role in vertebrate ion channel gating (36), the arrangement of TMIE C43 palmitoylation with respect to the TM6 π-helical turn suggests that could be important for gating of the ion channel.

## Discussion

The isolation and elucidation of the molecular structure of the native *C. elegans* TMC-2 complex, while similar to the TMC-1 complex, reveals several key features that may render it functionally distinct from TMC-1. First, the composition of the TMC-1 and TMC-2 complexes is not identical; the auxiliary subunit ARRD-6 does not co-purify with the TMC-2, which could be due to differences in expression pattern between TMC-1 and TMC-2. Second, TMC-1 and TMC-2 differ in the structure of the dimer interface, an observation that is intriguing due to the extremely high degree of structural conservation between TMC-1 and TMC-2 in all other regions of the proteins. Third, we observed key differences in the composition of pore-lining amino acids, 75% of which are localized to TM4, a pore-lining helix that is important for gating in vertebrate TMC-1.

Mass spectrometry and cryo-EM experiments of TMC-1 and TMC-2 provide further insight into the subunit composition of these complexes. Our results indicate that TMC-1 and TMC-2 do not form heterodimers in *C. elegans*, despite their conserved complex architecture and common localization to worm body wall and vulval muscles. Further, although worms express orthologs of the MT subunits protocadherin-15 (PCDH15) and lipoma HMGIC fusion partner-like 5 (LHFPL5), we did not observe co-purification of these subunits with worm TMC-1 and TMC-2. In vertebrates, PCDH15 forms part of the extracellular tip-link that connects adjacent stereocilia in a hair cell and is thought to convey force to the MT complex, thus playing an essential role in mechanosensation (37, 38). LHFPL5 forms a complex with PCDH15 *in vitro* (39) and null mutations in mouse LHFPL5 reduce MT currents (40). While it is possible that the lack of worm TMC-1/2 co-purification with PCDH15 or LHFPL5 is due to complex dissociation during purification, it is also conceivable that worms rely on a different mechanism of force-sensing that does not involve an extracellular tether. Finally, the presence of a π-helical turn in the likely pore-lining TMC-1/2 TM6 helix, and its interaction with the TMIE C43 palmitoyl group, may have implications for the gating mechanism of TMC-1/2 (Figure 5c). The π-helix structural motif is present in ∼15% of proteins and is typically associated with a specific protein function (41, 42), including ion channel gating (43, 44). We hypothesize that the π-helical turn plays role in mechanical gating of TMC-1/2, possibly by undergoing a π-to α-helix transition. In support of this speculation, the ion channels OSCA1.2 and TMEM16, both of which are evolutionarily related to TMC proteins, also harbor π-helical turns in their pore-forming helices. The mechanosensitive ion channel OSCA1.2 bears two π-helical turns in TM5 and TM6a (26), and the mouse Ca^2+^-activated chloride channel TMEM16A and the fungus lipid scramblase TMEM16 each harbor a π-helical turn in TM6 (31, 44). The gating mechanism of TMEM16A relies on an α-to-π transition, wherein Ca^2+^ binding stabilizes the open conformation of the channel via Ca^2+^ interaction with the π-helical turn (44). It is interesting to speculate that the TMC complexes have a similar gating mechanism. The interaction between the TMIE C43 palmitoyl group and hydrophobic residues on the TMC π-helical turn may stabilize the π-helix, thereby stabilizing the closed conformation of the channel. Deformation of the membrane through mechanical forces could disrupt interaction between the palmitoyl group and TM6, enabling a π-to-α transition that gates the ion channel.

Together, the conserved protein-lipid interactions between TMIE and TMC-1/2 allow us to build upon our proposed model of TMC complex mechanical activation (Figure 5c). The TMIE C43 and C44 palmitoyl groups interact with the pore-forming helices TM6 and TM8, as well as with the conserved ‘bridge’ phospholipid, which contacts both the TMC and TMIE transmembrane domains. These extensive lipid-mediated hydrophobic contacts may functionally link TMIE and TMC-1/2. We hypothesize that when force is applied to TMIE, either through membrane stretch or the action of an auxiliary subunit, those forces are translated to the TMC pore through lipid-mediated interactions, leading to channel opening and ion permeation.

## Materials and Methods

### Transgenic worm design

The strain PHX4776 *tmc-2(syb2106syb4776)* was generated by SunyBiotech using CRISPR/Cas9 genome editing and is referred to as the *tmc2::mVenus* line. A nucleic acid sequence encoding a mVenus-3xFLAG tag was inserted prior to the stop codon of the endogenous *tmc-2* gene (Wormbase: B0416.1a). The genotype was confirmed using PCR and primers ER14-seq-s (TGAACCGCATCGAGCTGA) and ER14-seq-a (CAAAAGTGGCCAATTTCGGG), which bind 104 bp upstream and 360 bp downstream from the insertion site, respectively, to amplify the region of interest. To enable elution of the engineered TMC-2 complex from affinity chromatography resin, a nucleic acid sequence coding for a PreScission protease (3C) cleavage site was placed between elements encoding the mVenus fluorophore and the 3xFLAG tag.

### Spectral confocal imaging

Adult worms were immobilized in M9 buffer (22 mM KH_2_PO_4_, 42 mM Na_2_HPO_4_, 86 mM NaCl, and 1 mM MgCl_2_) containing 30 mM sodium azide and placed on slides that were prepared with ∼4 mm agar pads. Spectral images were acquired on a Zeiss 34-channel LSM 880 Fast Airyscan inverted microscope with a 40x 1.2 NA water-immersion objective lens. Linear unmixing was employed to distinguish between the mVenus signal and autofluorescence. The autofluorescence signal was subtracted from each image. The 3D z-stack information is presented in 2D after performing a maximum intensity projection.

### Large scale *C. elegans* culture

Large-scale worm growth and optimization of TMC-2 protein isolation conditions were performed as described in Clark et al (23). Briefly, nematode growth medium (NGM) agar plates were prepared and spread with *E. coli* strain HB101, allowing the bacterial lawn to grow overnight at 37 °C. Worms were transferred to the NGM plates and grown for 3-4 days at 20 °C until HB101 cells were depleted. Worms on the plates were transferred to a liquid medium in 2 L baffled flasks, supplemented with HB101 (∼15 g per 500 mL medium) and streptomycin (50 μg/mL), and worms were grown at 20 °C with vigorous shaking (150 rpm) for 70-72 hours. To harvest worms, the liquid culture flasks were placed on ice for 1 hour to allow the worms to settle. The media was removed, and the worm slurry was collected in a tube, washed twice with 50 mL of ice cold M9 buffer by successive centrifugation (800 x g for 1 minute) and resuspended. Worms were ‘cleaned’ by sucrose density centrifugation at 1500 x g for 5 minutes after bringing the volume of worm slurry up to 25 mL with M9 buffer and adding 25 mL of ice cold 60% (w/v) sucrose. The worm layer on top was recovered and placed in a new tube and then washed once with 50 mL of ice cold M9 buffer and once with 50 mL of lysis buffer (50 mM Tris-Cl, pH 8.0, 50 mM NaCl, 5 mM EDTA, and protease inhibitors that included 0.8 μM aprotinin, 2 μg/mL leupeptin and 2 μM pepstatin). The volume of the worm pellet was measured and the same volume of lysis buffer was added to the tube and worm balls were made by dripping the slurry into liquid nitrogen. The worm balls were stored at −80 °C until further use.

### Isolation of the native TMC-2 complex

The experimental procedure for isolation of the TMC-2 complex is similar to the isolation procedure for the TMC-1 complex (22), but is nonetheless described in detail below. Approximately 50 g of frozen worm balls were disrupted using a ball mill (MM400, Retsch) where the grinding jar and ball were pre-cooled in liquid nitrogen. Disrupted worm powder was solubilized at 4 °C for 2 hours in a buffer containing 50 mM Tris-Cl (pH 8.0), 50 mM NaCl, 5 mM EDTA, 2% (w/v) glyco-diosgenin (GDN), and protease inhibitors (0.8 μM aprotinin, 2 μg/mL leupeptin and 2 μM pepstatin). The supernatant was clarified by centrifugation at 40,000 rpm (186,000 x g) for 50 minutes, followed by filtration with a 0.45 μm filter. The clarified supernatant was applied to anti-FLAG M2 magnetic affinity resin and incubated overnight on a rotator at 4°C. The resin was washed 5 times with a buffer containing 20 mM Tris-Cl (pH 8.0), 150 mM NaCl and 0.02% (w/v) GDN, using a volume of buffer that was 200-fold the volume of the resin. The TMC-2 complex was eluted by incubating the resin with 40 μg of 3C protease at 4 °C for 2 hours, on a rotator. Subsequently, the solution was supplemented with 1 mM CaCl_2_, final concentration, and the concentrate was loaded onto a size-exclusion chromatography (SEC) column (Superose 6 Increase 10/30 GL, GE Healthcare) connected to a Shimadzu high-pressure liquid chromatography instrument. The column was equilibrated in a buffer composed of 20 mM Tris-Cl (pH 8.0), 150 mM NaCl, 0.02% (w/v) GDN and 1 mM CaCl_2_ and the protein elution was monitored by tryptophan fluorescence (excitation= 280 nm, emission = 350 nm). The peak fractions from the putative dimeric TMC-2 complex were collected by hand, pooled, and concentrated for cryo-EM grid preparation. Approximately 100 ng of TMC-2 was isolated from 50 g of worm balls or about 3.8 × 10^7^ worms. The amount of protein was determined via mVenus fluorescence based on a standard plot. The isolated native TMC-2 sample was analyzed by SDS-PAGE (sodium dodecyl sulfate– polyacrylamide gel electrophoresis) and the protein bands were visualized by silver staining. For mass-spectrometry analysis, the putative dimeric TMC-2 complex peak was pooled and concentrated to a volume of 200 μL.

### Mass spectrometry

The purified TMC-2 complex sample was dried, dissolved in 5% sodium dodecyl sulfate, 8 M urea, 100 mM glycine (pH 7.55), reduced with (tris(2-carboxyethyl)phosphine (TCEP) at 37 °C for 15 min, alkylated with methyl methanethiosulfonate for 15 min at room temperature followed by addition of acidified 90% methanol and 100 mM triethylammonium bicarbonate buffer (TEAB; pH 7.55). The sample was then digested in an S-trap micro column briefly with 2 μg of a Tryp/LysC protease mixture, followed by a wash and 2 hr digestion at 47 °C with trypsin. The peptides were eluted with 50 mM TEAB and 50% acetonitrile, 0.2% formic acid, pooled and dried. Each sample was dissolved in 20 μL of 5% formic acid and injected into Thermo Fisher QExactive HF mass spectrometer. Protein digests were separated using liquid chromatography with a Dionex RSLC UHPLC system, then delivered to a QExactive HF (Thermo Fisher) using electrospray ionization with a Nano Flex Ion Spray Source (Thermo Fisher) fitted with a 20 μm stainless steel nano-bore emitter spray tip and 2.6 kV source voltage. Xcalibur version 4.0 was used to control the system. Samples were applied at 10 μL/min to a Symmetry C18 trap cartridge (Waters) for 10 min, then switched onto a 75 μm x 250 mm NanoAcquity BEH 130 C18 column with 1.7 μm particles (Waters) using mobile phases water (A) and acetonitrile (B) containing 0.1% formic acid, 7.5-30% acetonitrile gradient over 60 min and 300 nL/min flow rate. Survey mass spectra were acquired over m/z 375-1400 at 120,000 resolution (m/z 200) and data-dependent acquisition selected the top 10 most abundant precursor ions for tandem mass spectrometry by higher energy collisional dissociation using an isolation width of 1.2 m/z, normalized collision energy of 30 and a resolution of 30,000. Dynamic exclusion was set to auto, charge state for MS/MS +2 to +7, maximum ion time 100 ms, minimum AGC target of 3 × 10^6^ in MS1 mode and 5 × 10^3^ in MS2 mode. Data analysis was performed using Comet (v. 2016.01, rev. 3) (45) against a January 2022 version of canonical FASTA protein database containing *C. elegans* uniprot sequences and concatenated sequence-reversed entries to estimate error thresholds and 179 common contaminant sequences and their reversed forms. Comet searches for all samples performed with trypsin enzyme specificity with monoisotopic parent ion mass tolerance set to 1.25 Da and monoisotopic fragment ion mass tolerance set at 1.0005 Da. A static modification of +45.9877 Da was added to all cysteine residues and a variable modification of +15.9949 Da on methionine residues. A linear discriminant transformation was used to improve the identification sensitivity from the Comet analysis (46, 47). Separate histograms were created for matches to forward sequences and for matches to reversed sequences for all peptides of seven amino acids or longer. The score histograms of reversed matches were used to estimate peptide false discovery rates (FDR) and set score thresholds for each peptide class. The overall protein FDR was 1.2%.

### Cryo-EM sample preparation

A volume of 3.5 μL of the concentrated TMC-2 complex was applied to a Quantifoil grid (R2/1 300 gold mesh, covered by 2 nm continuous carbon film), which was glow-discharged at 15 mA for 30 seconds in the presence of amylamine. The grids were blotted and flash frozen using a Vitrobot mark IV for 2.5 seconds with 0 blot force after 30 seconds wait time under 100% humidity at 10 °C. The grids were plunge-frozen into liquid ethane, cooled by liquid nitrogen.

### Data acquisition

The native TMC-2 complex dataset was collected on a 300 keV FEI Titan Krios microscope equipped with a K3 detector. The micrographs were acquired in super-resolution mode (0.4155 Å/pixel) with a magnification of 105 kx corresponding to a physical pixel size of 0.831 Å/pixel. Images were collected by a 3×3 multi-hole per stage shift and a 6 multi-shot per hole method using Serial EM, with a defocus range of −1.5 to −2.5 μm. Each movie stack was exposed for 5.19 seconds and consisted of 50 frames per movie, with a total dose of 50 e^−^/Å^2^. A total of 23,364 movies were collected.

### Image processing

Beam-induced motion was corrected by patch motion correction with an output Fourier cropping factor of 1/2 (0.831 Å/pixel). Contrast transfer function (CTF) parameters were estimated by patch CTF estimation in CryoSparc v3.3.1 (48). A total of 23,324 movies were selected by manual curation and the particles were picked using blob-picker with minimum and maximum particle diameters of 140 Å and 200 Å, respectively (Figure S2a). Initially, 4.7 million particles were picked and extracted with a box size of 400 pixels and binned 4x (3.324 Å/pixel). 2D classification of all 4.7 million particles revealed that the TMC-2 complex exists in at least two assembly states, a ‘cis-dimer’ and a ‘trans-dimer’, as well as in two stoichiometric states, monomer and dimer (Figure S2b). As an initial step to remove TMC-2 complexes that were in the ‘trans-dimer’ or monomeric states, in addition to junk particles and empty micelles, five rounds of 2D classification were performed, resulting in 1.3 million particles in total. *Ab initio* reconstruction of the TMC-2 complex from the cleaned particle stack was unsuccessful, likely due to conformational heterogeneity. We expected the overall architecture of the TMC-2 complex to be similar to that of the TMC-1 complex for three reasons: (1) the 2D class averages of TMC-2 and TMC-1 are similar (Figure S3b and reference (22)) (2) TMC-1 and TMC-2 share 59% sequence identity in their transmembrane domains and (3) mass spectrometry analysis of the TMC-2 complex suggests that TMC-2 interacts with CALM-1 and TMIE, the same auxiliary proteins present in the TMC-1 complex. As starting models for heterogeneous refinement, we therefore used the ‘expanded’ form of the TMC-1 complex (PDB 7USW) at 3.0 Å resolution and the TMC-1 complex that had been low pass filtered to 10 Å and 20 Å. The full particle stack consisting of 1.3 million particles from 2D classification was subjected to heterogeneous refinement with C1 symmetry using these three starting models in six classes total. A second round of heterogeneous refinement was then performed with the two good classes, which consisted of 623k particles, to further clean the particle stack. The best class, consisting of 320k particles, was selected for a final round of heterogeneous refinement, yielding 213k particles that were re-extracted from unbinned images. To attain higher resolution and improve map quality, non-uniform refinement, including defocus and global CTF refinement, was performed in Cryosparc v3.3.1, resulting in a map with 3.0 Å resolution. One protomer of this map was clearly lower resolution than the other, likely due to conformational heterogeneity of the dimer interface. To further improve the resolution of one protomer, local refinement was performed with a mask covering one half of the complex (one subunit each of TMC-2, CALM-1 and TMIE). This resulted in the ‘single protomer map’ with a resolution of 2.93 Å.

The ‘protomer’ map lacked adequate resolution of TM10 for model building. To attain higher resolution in this region, the re-extracted particles were first symmetry expanded with C2 symmetry. The particles were then subjected to 3D classification without alignment using a mask centered on TM10 with 6 Å target resolution, followed by local refinement. The resulting ‘dimer interface’ map was used for model building of TMC-2 TM10.

### Structure determination and model building

The ‘protomer’ map was used for building the entirety of the TMC-2 complex structure aside from TM10 (Figure S2a). The transmembrane helices of TMC-2 (TM1-9, excluding TM10), predicted by Alphafold2 (49) as a template, were fit into the map with rigid body fitting and *de novo* model building in Coot (50) and Isolde (51). The cytoplasmic and extracellular regions of TMC-2 were built with a combination of *de novo* model building and rigid body fitting of the TMC-1 structure (PDB 7USW), followed by appropriate modification of the amino acid sequence. The possible ion permeation pore of the channel was determined by MOLE 2.5 (52). Carbohydrate groups were modeled into protruding densities of N225 and N748 on TMC-2, at predicted *N*-linked glycosylation sites. To build the structures of the auxiliary subunits CALM-1 and TMIE, the subunit structures from the TMC-1 complex were fit into the map with rigid body fitting in UCSF Chimera. The TMIE and CALM-1 subunits were then manually adjusted in Coot, followed by real-space refinement in Phenix.

As the ‘protomer’ map only includes density for one half of the TMC-2 complex, it was necessary to duplicate the map to build the second protomer. The map was rotated 180 degrees and rigid body fit into the ‘dimer interface’ map in UCSF Chimera (53) to produce a map with two copies of each subunit. The model of the TMC-2 complex was then rigid body fit into the map in UCSF Chimera, followed by manual adjustment in Coot. The TM10 helices were built *de novo* in Coot using the ‘dimer interface’ map (Figure S2a). The model was refined against the sharpened ‘protomer’ map by real-space refinement in Phenix.

The following regions of TMC-2 were not modeled into the map because of weak or absent densities: The N-terminal region of TMC-1 (M1 to A88), the predicted loop region between TM5 and TM6 (A472 to T582) and the C-terminal region (M814 to D1203). The side chains with weak density located in the TM10 helices were modeled as alanine residues. The N-terminal region of CALM-1, from residues M1 to F14, and the N- and C-terminal residues of TMIE, M1-A13 and K63-V117, were not modeled due to a lack of density.

The palmitoylation of TMIE C43 in the TMC-1 complex was built using *de novo* model building in Coot, based on the cryo-EM volume data previously deposited to the Electron Microscopy Data Bank (22). Real space refinement and regularization were performed in Coot.

## Acknowledgments

We thank A. Aballay, F. Jalali-Yazdi, and members of the Gouaux and Baconguis laboratories for helpful discussions; C. Sun for computer support; A. Chinn and M. Frisbie for help with worm growth; and R. Courtney for proof reading. We acknowledge the help of the staff in the OHSU Advanced Light Microscopy Core (RRID:SCR_009961), especially S. Petrie and B. Jenkins for help with worm spectral imaging. Mass spectrometric analysis was performed by the OHSU Proteomics Shared Resource with partial support from NIH core grants P30EY010572, P30CA069533, and OHSU Emerging Technology Fund. Initial cryo-EM grids were screened at the OHSU Multiscale Microscopy Core (MMC). The large single-particle dataset was collected at the Janelia Research Campus of the Howard Hughes Medical Institute (HHMI) with the help of S. Yang. This work was supported by NIH grant 1F32DC017894 to S.C. E.G. gratefully acknowledges J. LaCroute and B. LaCroute for support, and is an investigator of the HHMI.

## Data Availability

The volumes for the cryo-EM data have been deposited in the Electron Microscopy Data Bank under accession code EMD-41356 (‘protomer’ map) and EMD-41432 (‘dimer interface’ map). The coordinates have been deposited in the Protein Data Bank under accession code 8TKP. The TMC-1 complex coordinates (7USW, 7USX, and 7USY) have been revised to include the palmitoylation of TMIE C43.

## References

1. J. M. Kefauver, A. B. Ward, A. Patapoutian, Discoveries in structure and physiology of mechanically activated ion channels. Nature 587, 567–576 (2020).

2. B. Pan et al., TMC1 and TMC2 are components of the mechanotransduction channel in hair cells of the mammalian inner ear. Neuron 79, 504–515 (2013).

3. K. Kurima et al., TMC1 and TMC2 Localize at the Site of Mechanotransduction in Mammalian Inner Ear Hair Cell Stereocilia. Cell reports 12, 1606–1617 (2015).

4. S. W. Chou et al., A molecular basis for water motion detection by the mechanosensory lateral line of zebrafish. Nature communications 8, 2234 (2017).

5. I. V. Pacentine, T. Nicolson, Subunits of the mechano-electrical transduction channel, Tmc1/2b, require Tmie to localize in zebrafish sensory hair cells. PLoS Genet 15, e1007635 (2019).

6. T. Erickson et al., Integration of Tmc1/2 into the mechanotransduction complex in zebrafish hair cells is regulated by Transmembrane O-methyltransferase (Tomt). eLife 6 (2017).

7. Y. V. Zhang, T. J. Aikin, Z. Li, C. Montell, The Basis of Food Texture Sensation in Drosophila. Neuron 91, 863–877 (2016).

8. Y. Q. Tang et al., Ankyrin Is An Intracellular Tether for TMC Mechanotransduction Channels. Neuron 107, 759–761 (2020).

9. L. F. Corns, J. Y. Jeng, G. P. Richardson, C. J. Kros, W. Marcotti, TMC2 Modifies Permeation Properties of the Mechanoelectrical Transducer Channel in Early Postnatal Mouse Cochlear Outer Hair Cells. Front Mol Neurosci 10, 326 (2017).

10. G. Keresztes, H. Mutai, S. Heller, TMC and EVER genes belong to a larger novel family, the TMC gene family encoding transmembrane proteins. BMC Genomics 4, 24 (2003).

11. K. Kurima, Y. Yang, K. Sorber, A. J. Griffith, Characterization of the transmembrane channel-like (TMC) gene family: functional clues from hearing loss and epidermodysplasia verruciformis. Genomics 82, 300–308 (2003).

12. Y. Kawashima, K. Kurima, B. Pan, A. J. Griffith, J. R. Holt, Transmembrane channel-like (TMC) genes are required for auditory and vestibular mechanosensation. Pflugers Archiv: European journal of physiology 467, 85–94 (2015).

13. B. Pan et al., TMC1 Forms the Pore of Mechanosensory Transduction Channels in Vertebrate Inner Ear Hair Cells. Neuron 99, 736–753 e736 (2018).

14. A. Antonsson et al., Variants of EVER1 and EVER2 (TMC6 and TMC8) and human papillomavirus status in patients with mucosal squamous cell carcinoma of the head and neck. Cancer Causes Control 27, 809–815 (2016).

15. J. Song et al., Pan-Cancer Analysis Reveals the Signature of TMC Family of Genes as a Promising Biomarker for Prognosis and Immunotherapeutic Response. Front Immunol 12, 715508 (2021).

16. Y. Guo et al., Transmembrane channel-like (tmc) gene regulates Drosophila larval locomotion. Proceedings of the National Academy of Sciences of the United States of America 113, 7243–7248 (2016).

17. J. Dao, A. Lee, D. K. Drecksel, N. M. Bittlingmaier, T. M. Nelson, Characterization of TMC-1 in C. elegans sodium chemotaxis and sodium conditioned aversion. BMC Genet 21, 37 (2020).

18. X. Wang, G. Li, J. Liu, J. Liu, X. Z. Xu, TMC-1 Mediates Alkaline Sensation in C. elegans through Nociceptive Neurons. Neuron 91, 146–154 (2016).

19. X. Yue et al., TMC Proteins Modulate Egg Laying and Membrane Excitability through a Background Leak Conductance in C. elegans. Neuron 97, 571–585 e575 (2018).

20. L. Zhang et al., TMC-1 attenuates C. elegans development and sexual behaviour in a chemically defined food environment. Nature communications 6, 6345 (2015).

21. J. Wu et al., GABA signaling triggered by TMC-1/Tmc delays neuronal aging by inhibiting the PKC pathway in C. elegans. Sci Adv 8, eadc9236 (2022).

22. H. Jeong et al., Structures of the TMC-1 complex illuminate mechanosensory transduction. Nature 610, 796–803 (2022).

23. S. Clark, H. Jeong, A. Goehring, Y. Kang, E. Gouaux, Large-scale growth of C. elegans and isolation of membrane protein complexes. Nat Protoc 10.1038/s41596-023-00852-5 (2023).

24. T. Kawate, E. Gouaux, Fluorescence-detection size-exclusion chromatography for precrystallization screening of integral membrane proteins. Structure 14, 673–681 (2006).

25. X. Bai et al., Caenorhabditis elegans PIEZO channel coordinates multiple reproductive tissues to govern ovulation. eLife 9 (2020).

26. S. Jojoa-Cruz et al., Cryo-EM structure of the mechanically activated ion channel OSCA1.2. eLife 7 (2018).

27. K. Saotome et al., Structure of the mechanically activated ion channel Piezo1. Nature 554, 481–486 (2018).

28. A. P. J. Giese et al., CIB2 interacts with TMC1 and TMC2 and is essential for mechanotransduction in auditory hair cells. Nature communications 8, 43 (2017).

29. B. Zhao et al., TMIE is an essential component of the mechanotransduction machinery of cochlear hair cells. Neuron 84, 954–967 (2014).

30. C. L. Cunningham et al., TMIE Defines Pore and Gating Properties of the Mechanotransduction Channel of Mammalian Cochlear Hair Cells. Neuron 107, 126–143 e128 (2020).

31. J. D. Brunner, N. K. Lim, S. Schenck, A. Duerst, R. Dutzler, X-ray structure of a calcium-activated TMEM16 lipid scramblase. Nature 516, 207–212 (2014).

32. S. Dang et al., Cryo-EM structures of the TMEM16A calcium-activated chloride channel. Nature 552, 426–429 (2017).

33. M. Zhang et al., Structure of the mechanosensitive OSCA channels. Nat Struct Mol Biol 25, 850–858 (2018).

34. R. Fettiplace, K. X. Kim, The physiology of mechanoelectrical transduction channels in hearing. Physiol Rev 94, 951–986 (2014).

35. M. Beurg, A. Barlow, D. N. Furness, R. Fettiplace, A Tmc1 mutation reduces calcium permeability and expression of mechanoelectrical transduction channels in cochlear hair cells. Proceedings of the National Academy of Sciences of the United States of America 116, 20743–20749 (2019).

36. N. Akyuz et al., Mechanical gating of the auditory transduction channel TMC1 involves the fourth and sixth transmembrane helices. Sci Adv 8, eabo1126 (2022).

37. P. Kazmierczak et al., Cadherin 23 and protocadherin 15 interact to form tip-link filaments in sensory hair cells. Nature 449, 87–91 (2007).

38. H. Sakaguchi, J. Tokita, U. Muller, B. Kachar, Tip links in hair cells: molecular composition and role in hearing loss. Curr Opin Otolaryngol Head Neck Surg 17, 388–393 (2009).

39. J. E. Ge, J.; Goehring, A; Zhao, J.; Schuck, P.; Gouaux E., Structure of mouse protocadherin 15 of the stereocilia tip link in complex with LHFPL5. eLife 7 (2018).

40. M. Beurg, W. Xiong, B. Zhao, U. Muller, R. Fettiplace, Subunit determination of the conductance of hair-cell mechanotransducer channels. Proceedings of the National Academy of Sciences of the United States of America 112, 1589–1594 (2015).

41. K. R. Rajashankar, S. Ramakumar, Pi-turns in proteins and peptides: Classification, conformation, occurrence, hydration and sequence. Protein Sci 5, 932–946 (1996).

42. R. B. Cooley, D. J. Arp, P. A. Karplus, Evolutionary origin of a secondary structure: pi-helices as cryptic but widespread insertional variations of alpha-helices that enhance protein functionality. J Mol Biol 404, 232–246 (2010).

43. L. Zubcevic, S. Y. Lee, The role of pi-helices in TRP channel gating. Curr Opin Struct Biol 58, 314–323 (2019).

44. C. Paulino, V. Kalienkova, A. K. M. Lam, Y. Neldner, R. Dutzler, Activation mechanism of the calcium-activated chloride channel TMEM16A revealed by cryo-EM. Nature 552, 421–425 (2017).

45. J. K. Eng, T. A. Jahan, M. R. Hoopmann, Comet: an open-source MS/MS sequence database search tool. Proteomics 13, 22–24 (2013).

46. P. A. Wilmarth, M. A. Riviere, L. L. David, Techniques for accurate protein identification in shotgun proteomic studies of human, mouse, bovine, and chicken lenses. J Ocul Biol Dis Infor 2, 223–234 (2009).

47. A. Keller, A. I. Nesvizhskii, E. Kolker, R. Aebersold, Empirical statistical model to estimate the accuracy of peptide identifications made by MS/MS and database search. Anal Chem 74, 5383–5392 (2002).

48. A. Punjani, J. L. Rubinstein, D. J. Fleet, M. A. Brubaker, cryoSPARC: algorithms for rapid unsupervised cryo-EM structure determination. Nature methods 14, 290–296 (2017).

49. J. Jumper et al., Highly accurate protein structure prediction with AlphaFold. Nature 596, 583–589 (2021).

50. P. Emsley, B. Lohkamp, W. G. Scott, K. Cowtan, Features and development of Coot. Acta Crystallogr D Biol Crystallogr 66, 486–501 (2010).

51. T. I. Croll, ISOLDE: a physically realistic environment for model building into low-resolution electron-density maps. Acta Crystallogr D Struct Biol 74, 519–530 (2018).

52. L. Pravda et al., MOLEonline: a web-based tool for analyzing channels, tunnels and pores (2018 update). Nucleic Acids Res 46, W368–W373 (2018).

53. E. F. Pettersen et al., UCSF Chimera--a visualization system for exploratory research and analysis. J Comput Chem 25, 1605–1612 (2004).

