## Supplementary Information for "Structure of *C. elegans* TMC-2 complex suggests roles of lipid-mediated subunit contacts in mechanosensory transduction"

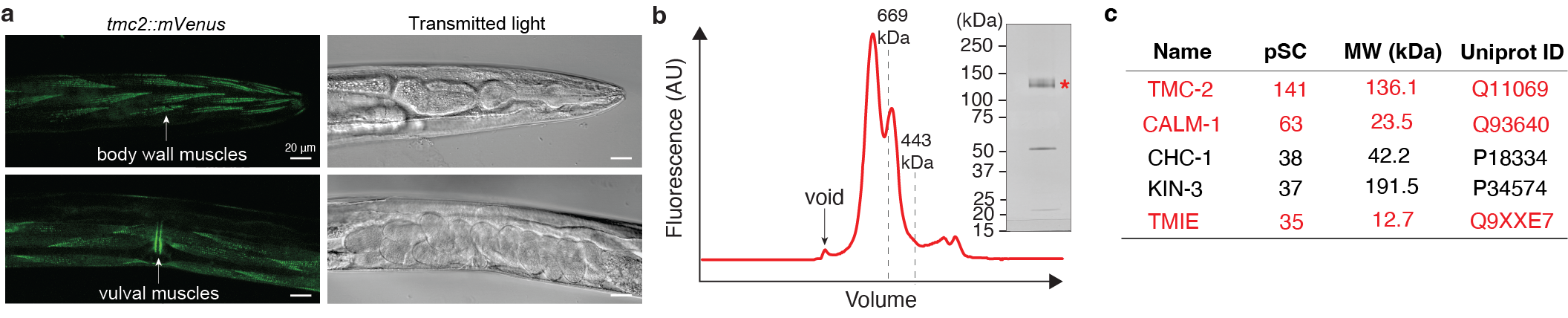

**Fig. S1: Isolation of TMC-2 from *C. elegans***. **a,** Spectral confocal images of mVenus fluorescence and transmitted light in a *tmc-2::mVenus* worm showing fluorescence in the body wall and vulval muscles. **b,** Representative FSEC profile of the TMC-2 complex, detected via mVenus fluorescence. Inset shows a silver stained SDS-PAGE gel of the purified TMC-2 complex. Red asterisk indicates TMC-2. **c,** Mass spectrometry analysis of the purified TMC-2 complex, showing the five proteins with the highest peptide spectral counts (pSC). Components of the TMC-2 complex that were also identified in the TMC-2 single particle cryo-EM structure are highlighted in red.

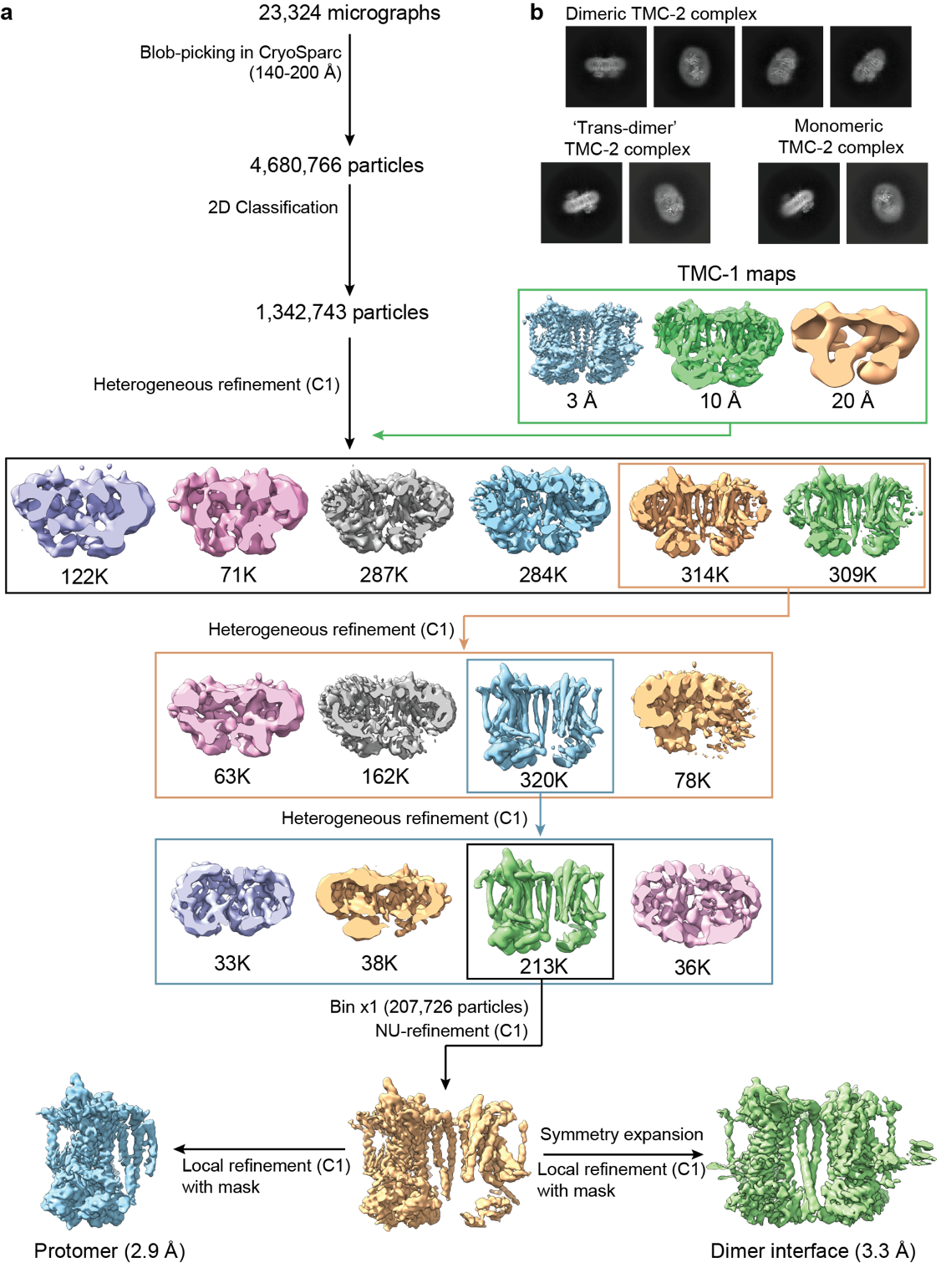

**Fig. S2: Cryo-EM data analysis of the TMC-2 complex. a,** Flow chart of cryo-EM data analysis of the TMC-2 complex. **b,** Representative 2D classes of the dimer TMC-2 complex (top) as well as the ‘trans-dimer’ and monomeric classes, the latter two of which were discarded during heterogeneous refinement.

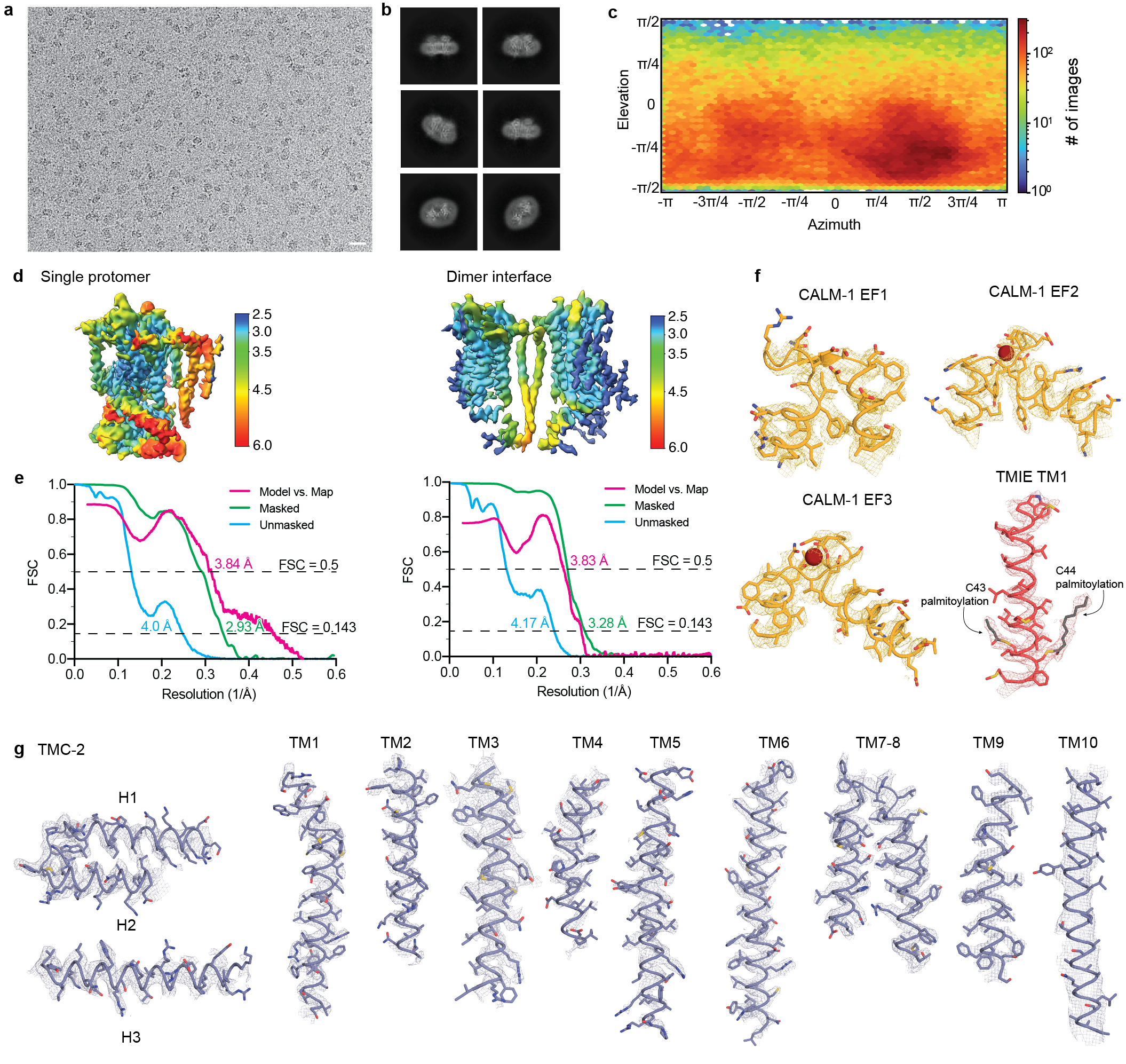

**Fig. S3: Cryo-EM classes, statistics, angular distributions and selected sections of density maps. a,** A representative cryo-EM micrograph of the TMC-2 particles. Scale bar = 200 Å. **b,** Selected 2D classes of the TMC-2 complex. **c,** Angular distributions of the final reconstruction. **d,** Density maps colored by local resolution values. **e,** Fourier shell correlations (FSEC) curves for each map and model. **f,** Fragments of cryo-EM density map and atomic models of the auxiliary subunits, CALM-1 and TMIE. Cryo-EM maps are shown as yellow and red mesh. **g,** Fragments of cryo-EM density map and atomic models of TMC-2. Cryo-EM maps are shown as purple mesh.

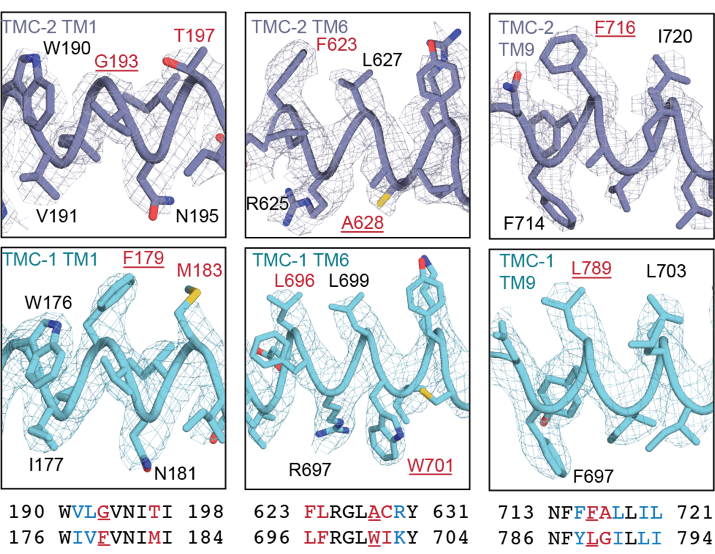

**Fig. S4.** Density feature differences between the TMC-1 and TMC-2 maps. The TMC-2 map and model are shown in purple and the TMC-1 map and model are shown in cyan. A sequence alignment for each fragment is shown at the bottom of the panel with the same coloring as in panel (d) of Figure 2. Underlined residues are non-conservative substitutions that are clearly distinguished in the density maps. The sequence of TMC-2 is on top and the sequence of TMC-1 is on bottom.

**Table S1.** Statistics for 3D reconstruction and model building

| Codes | PDB 8TKP |
| --- | --- |
|  | Protomer: EMD-41356  Dimer interface: EMD-41432 |
| **Data collection and processing** | |
| Microscope | Titan Krios |
| Camera | K3 BioQuantum |
| Magnification | 105,000 |
| Voltage (kV) | 300 |
| Defocus range (μm) | -1.5 to -2.5 |
| Exposure time (s) | 5.19 |
| Dose rate (*e^-^*/Å^2^/s) | 7.98 |
| Number of frames | 50 |
| Pixel size (Å) | 0.831 (0.4155 super-resolution) |
| Micrographs (no.) | 23,364 |
| Symmetry imposed | C1 |
| Initial particles (no.) | 4,680,766 |
| Final particles (no.) | 207,726 |
| Map resolution (Å) | 2.9 (protomer), 3.3 (dimer interface) |
| FSC threshold | 0.143 |
| **Refinement** | |
| Initial model (PDB code) | Alphafold, 7USW |
| Model resolution (Å) | 3.2 |
| FSC threshold | 0.5 |
| Model composition |  |
| Non-hydrogen atoms | 14666 |
| Protein atoms | 13848 |
| Ligand atoms | 818 |
| *B* factors (Å^2^) |  |
| Protein | 45.38 |
| Lipids and calcium | 66.06 |
| R.m.s. deviations |  |
| Bond length (Å) | 0.002 |
| Bond angle (˚) | 0.540 |
| **Validation** | |
| Favored (%) | 98.04 |
| Allowed (%) | 1.96 |
| Disallowed (%) | 0 |
| Poor rotamers (%) | 1.07 |
| MolProbity score | 1.34 |
| Clash score | 5.79 |
